# Structure and mechanism of human diacylglycerol acyltransferase 1

**DOI:** 10.1101/2020.01.06.896332

**Authors:** Lie Wang, Hongwu Qian, Yin Nian, Yimo Han, Zhenning Ren, Hanzhi Zhang, Liya Hu, B. V. Venkataram Prasad, Nieng Yan, Ming Zhou

**Author notes:** These authors contributed equally (Lie Wang, Hongwu Qian, Yin Nian, Yimo Han). Correspondence to Ming Zhou and Nieng Yan.

## Abstract

Human diacylglycerol O-acyltransferase-1 (hDGAT1) synthesizes triacylglycerides and is required for dietary fat absorption and fat storage. The lack of 3-dimensional structure has limited our understanding of substrate recognition and mechanism of catalysis, and hampers rational targeting of hDGAT1 for therapeutic purposes. Here we present the structure of hDGAT1 in complex with a substrate oleoyl Coenzyme A at 3.1 Å resolution. hDGAT1 forms a homodimer and each protomer has nine transmembrane helices that carve out a hollow chamber in the lipid bilayer. The chamber encloses highly conserved catalytic residues and has separate entrances for the two substrates fatty acyl Coenzyme A and diacylglycerol. The N-terminus of hDGAT1 makes extensive interactions with the neighboring protomer, and is required for enzymatic activity.

## Introduction

Diacylglycerol acyltransferase 1 (DGAT1, EC 2.3.1.20) is an integral membrane protein that synthesizes triacylglycerides (TG or triglycerides, Figure S1) from two substrates, diacylglycerol (DAG or diglyceride) and fatty acyl Coenzyme A (acyl-CoA) (1). The primary physiological function of DGAT1 in mammals is fat absorption and storage (2, 3). In humans, DGAT1 is highly expressed in epithelial cells of the small intestine and DGAT1 activity is essential for dietary fat absorption (4, 5). DGAT1 is also found in other organs or tissues such as liver where it synthesizes fat for storage and in female mammary glands where it produces fat in the milk (6). In plants, DGAT1 synthesizes seed storage lipids (seed oils) that are widely utilized for food or biofuels (7, 8). In addition to the synthesis of TG, hDGAT1 is also implicated as a host virulence factor for hepatitis C and rotaviruses (9–11). *Dgat1*^-/-^ mice are viable and show significantly reduced TG in all tissues, increased sensitivity to leptin and insulin, and resistance to obesity when kept on a high-fat diet (12, 13). These results have generated considerable interest in targeting hDGAT1 for treating hypertriglyceridemia and fatty liver disease, and for controlling obesity, diabetes, coronary heart diseases (14, 15).

DGAT1 belongs to a large superfamily of membrane-bound O-acyl transferases (MBOAT, http://pfam.xfam.org/family/MBOAT) that are found in all kingdoms of life. MBOAT family includes enzymes such as acyl-CoA:cholesterol acyltransferase (ACAT) that attaches a fatty acid to a cholesterol and is crucial for bile acid sysnthesis (16), and protein-serine O-palmitoleoyltransferase (PORCUPINE or PORCN) that adds a palmitoleate to a conserved serine residue to activate the WNT protein (17, 18). Members of the MBOAT family have a highly conserved histidine residue required for the transferase activity and are predicted to have 8-11 transmembrane segments (16, 19–24). Crystal structure of a bacterial member of MBOAT, DltB, was reported recently (25). DltB catalyzes the transfer of a D-alanine from an intracellular protein DltC to a lipoteichoic acid located on the extracellular side of the membrane (Figure S1). However, the structure of DltB is not a suitable model for hDGAT1 because of their low sequence identity (∼20%) and very different substrates. As we will see in Discussion, DltB and DGAT1 likely evolved from a common ancestor but have diverged significantly to accommodate differences in their substrates.

Because of the physiological and pharmacological importance of DGAT1 and its significance as a model for other related MBOAT members, it is important to understand the molecular details of how DGAT1 works in terms of substrate recognition and mechanism of catalysis and inhibition. We expressed and purified human DGAT1, and solved the structure of hDGAT1 in complex with a substrate oleoyl-CoA at an overall resolution of 3.1 Å by using single-particle cryo-electron microscopy (cryoEM).

## Functional characterization of purified hDGAT1

Full-length hDGAT1 was over-expressed and purified (Figure 1A, Methods). The purified hDGAT1 elutes on a size-exclusion column with a main peak at an elution volume of ∼11.7 ml, which corresponds to a molecular weight of ∼150 kDa (Figure S2A). Since each hDGAT1 protomer has a molecular weight of ∼55 kDa and with the detergent micelle, the elution peak is consistent with a dimeric hDGAT1. The dimer is also clearly visible on an SDS-PAGE, indicating that it is stable enough to be partially resistant to denaturing conditions (Figure 1A). We noticed that there is a minor peak at ∼10.4 ml that likely corresponds to a tetramer (Figures 1A and S2A). A number of previous studies also showed that DGAT1 from plants and mammals could form either dimer or tetramer (20, 26, 27), however, it is not clear whether both of the oligomeric states exist in the native endoplasmic reticulum membranes and whether the oligomeric state has an impact on the enzymatic functions.

**Figure 1.**
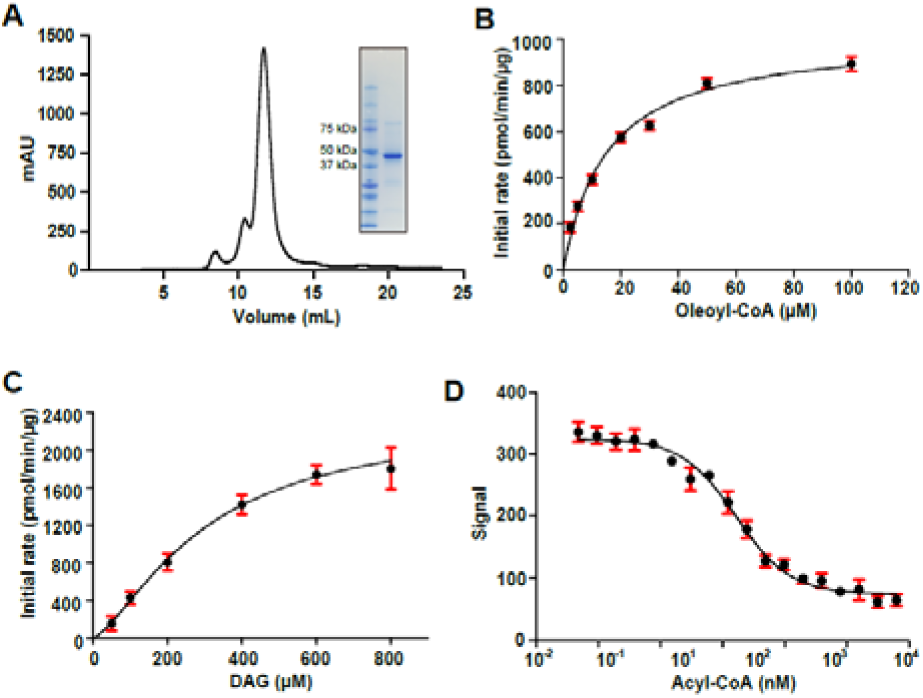
Purification and functional characterization of hDGAT1. **A**, Size-exclusion chromatography profile of purified hDGAT1. Inset: SDS–PAGE of the purified hDGAT1. **B-C.** Initial rate of reaction versus oleoyl-CoA (**B**) or DAG (**C**) concentration. Data in **B** were fit with a Michaelis-Menten equation, and data in **C** were fit with an allosteric sigmoidal equation (Methods). **D.** Competitive binding of oleoyl-CoA measured against its competition against 0.25 μM of ^3^H-acetyl-CoA. Data were fit with a single-site competitive binding isotherm (Methods). In **B**, **C** and **D**, Each symbol is the average of three repeats. Error bars are standard errors of the mean (s.e.m.)

During the purification process, a thick white layer of fat appeared after the centrifugation step (Figure S2B), indicating that the heterologously expressed hDGAT1 is functional in cells. To find out whether the purified hDGAT1 dimer remains functional, we measured the activity of the purified hDGAT1 by following the production of Coenzyme A (Figure S2C). Rapid production of Coenzyme A occurs in the presence of oleoyl-CoA, 1,2-dioleoyl-sn-glycerol (1,2-DAG) and the purified hDGAT1 (Figures S2C-S2D). In contrast, Coenzyme A production was not observed when either 1,2-DAG or hDGAT1 was omitted from the reaction mixtures. Coenzyme A production was almost completely suppressed in the presence of T863, a known hDGAT1 inhibitor (28) (Figure S2D). These results indicate that the purified dimeric hDGAT1 preserves its enzymatic function.

We further characterized the purified hDGAT1 to establish basic parameters of its enzymatic reaction. The initial rate of Coenzyme A production at different concentrations of oleoyl-CoA was measured and it follows a Michaelis-Menten type relationship with K_M_ and V_max_ of 14.6 ± 1.3 µM and 956.6 ± 36.1 nmol/mg/min, respectively (Figure 1). The K_M_ and V_max_ values are comparable to those previously reported of hDGAT1 in microsomes (29, 30), and the V_max_ value is equivalent to a turnover rate of ∼1/second for each hDGAT1 protomer. Enzymatic activity was also measured at different concentrations of DG which has a K_M_ and V_max_ of 597.1 ± 94.5 µM and 3310 ± 279.1 nmol/mg/min (Figure 1C). We did not observe substrate inhibition up to 100 μM of oleoyl-CoA, which is different from a previous study on DGAT1 from the plant μ

*Brassica napus* (31). hDGAT1 has almost no preference for oleoyl-CoA (K_M_= 14.6 ± 1.3 µM; V_max_= 956.6 ± 36.1 nmol/mg/min) versus stearoyl-CoA (K_M_= 8.6 ± 1.3 µM; V_max_= 839.4 ± 49.9 nmol/mg/min), or palmitoleoyl-CoA (K_M_= 6.2 ± 0.9 µM; V_max_ = 838.6 ± 31.6 nmol/mg/min) versus palmitoyl-CoA (K_M_ = 6.4 ± 1.1 µM; V_max_ = 767.8 ± 34.0 nmol/mg/min) as acyl donors but has a slower V_max_ and higher K_M_ for decanoyl-CoA (Figures S2G-S2I, Table S2). For acyl acceptors, hDGAT1 has a clear preference for DAG over the two mono-acyl glycerols (Figure S2F, Table S2). Divalent cations such as Mg^2+^ or Ca^2+^ has no significant effect on the enzymatic reaction (Figure S2E, Table S2), consistent with previous results from both the mammalian and plant DGAT1s (32–34). In addition to the enzymatic activity, we estimated binding affinity of oleoyl-CoA to hDGAT1 in a scintillation proximity assay in which we used oleoyl-CoA to compete the binding of ^3^H-acetyl-CoA, and the result shows that oleoyl-CoA binds to hDGAT1 with an IC50 of 18.2 ± 5.2 nM (Figure 1D).

## Overall structure of hDGAT1

hDGAT1 structure was solved by single-particle cryoEM. Due to the modest size of hDGAT1 particles (∼110 kDa), conditions for grid preparation and data collection were extensively optimized to achieve desired contrast and particle density (Figure S3A, Methods). Data were processed following the flow chart shown in Figure S3B and detailed in Methods. A density map was reconstructed to an overall 3.1 Å resolution with C2 symmetry imposed using 408,945 particles (Figure S3B). Furthur refinement using 275,945 particles produced a map that has almost identical resolution but better density for bound lipids and detergents (Figure S3B).

Resolution for helices close to the core of the dimer reaches 2.7 Å while regions close to the peripheral of the dimer has lower resolution likely due to their relatively higher mobility (Figure S3C).

The density map is of sufficient quality to allow *de novo* building of residues 64 to 224 and 239 to 481, which include all the transmembrane helices, one oleoyl-CoA, and 5 partially resolved lipid/detergent molecules, and the structure was refined to proper geometry (Figure S4, Table S1). The first 63 and the last 5 residues, and residues 225-238 which is part of a cytosolic loop, were not resolved. Residues 112 to 120, which is part of a luminal loop, were partially resolved and built as poly-alanines.

hDGAT1 dimer has a dimension of ∼105 by 55 by 48 Å and is shaped like a canoe (Figures. 2A-2D). Based on the positive-inside rule (35), the N-terminus of hDGAT1 resides at the cytosolic side (Figure S5). This assignment is also consistent with the previous consensus based on biochemical studies (20, 36, 37) and allows for unambiguous placement of the C-terminus to the lumen side of the ER. Each hDGAT1 protomer has nine transmembrane helices, TM1-9, and three long loops, an ER luminal (extracellular) loop EL1 between TM1 and 2, an intracellular loop IL1 between TM4 and 5, and a second intracellular loop IL2 between TM6 and 7 (Figures 2E-F). Both IL1 and IL2 are structured and composed of highly conserved amino acid sequences (Figure S6).

**Figure 2.**
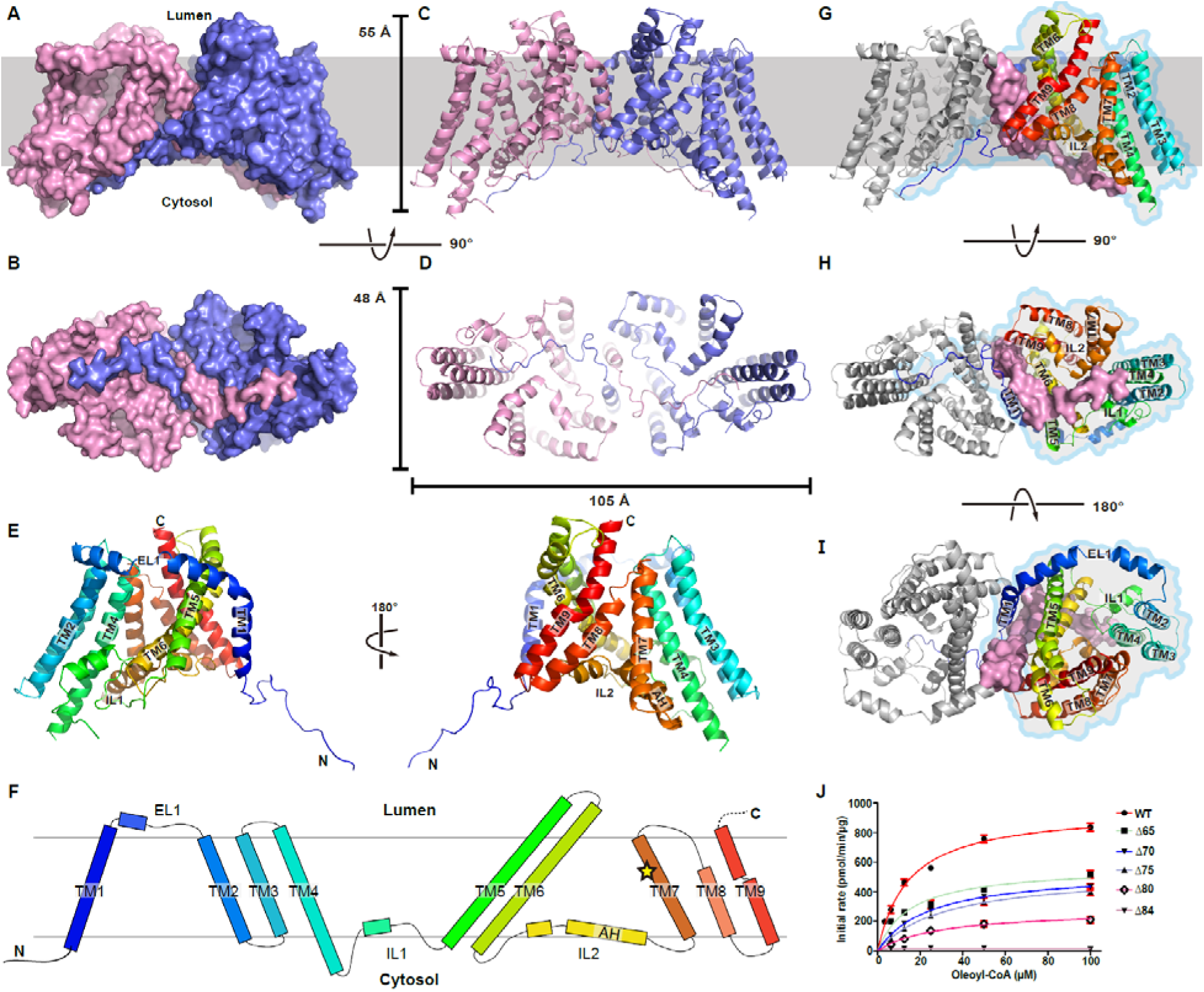
Structure of hDGAT1. **A**–**D** Structure of hDGAT1 dimer is shown in cartoon and surface representations as viewed from within the plane of the membrane (**A, C**), or the intracellular side of the membrane (**B, D**). Approximate position of the ER membrane is marked as grey shade. **E.** Cartoon representation of an hDGAT1 protomer in two orientations. **F**. Topology of hDGAT1. The position of His415 is marked as a yellow star. **G**-**I**, dimerization interface of hDGAT1 viewed in three orientations. One protomer is shown as grey cartoon but with its TM1 and the N-terminus in surface. The other protomer is shown as rainbow colored cartoon and marked with an outline. **J.** Enzymatic activity of N-terminal truncations of hDGAT1. Initial rate of reaction versus oleoyl-CoA concentration. Each symbol represents the average of three repeats. Error bars are s.e.m.. Solid lines are fit of the data points with a Michaelis-Menten equation.

In each protomer, TM2-9 and the two intracellular loops IL1 and 2 form a distinctive structural fold that we define as the MBOAT fold (Figures 2E-2F and and 3A-3D). TM1, which is not part of the MBOAT fold, is isolated from the rest of the transmembrane helices and linked to the MBOAT fold by the long ER luminal loop EL1 (residues 110 to 125). EL1 is partially structured and extends ∼35 Å along the luminal side of the protein (Figures 2E and 2I).

## The Dimer interface

Although TM1 seems suspended in the membrane when a protomer is viewed in isolation, the space between the TM1 and the rest of the protomer (the MBOAT fold) is filled by the TM1 from the neighboring protomer so that the two protomers form a domain-swapped homodimer (Figures 2A-F). TM1 makes extensive hydrophobic interactions with both TM6 and TM9 from the neighboring subunit (Figures 2G-I). Crossover of the TM1 helix brings the N-terminus of one protomer close to the intracellular side of its neighbor. Residues 64 to 80 of the N-terminus interact with both IL1 and IL2 of the neighboring subunit. Overall, the dimer interface has an extensive buried surface area of 684.5 Å^2^. The two TM1s only make a single contact at Ile80 located close to the intracellular side of the membrane, and the space between them is filled with 2 lauryl maltose neopentyl glycol (LMNG) molecules and 4 partially resolved lipid molecules (Figures S5A-H).

Previous studies on a plant DGAT1 have identified part of the N-terminus as intrinsically disordered protein, and showed that deletion of the N-terminus before TM1 led to a loss of the enzymatic activity (31, 38, 39). In the hDGAT1 structure, the first 63 residues are not resolved and likely is disordered while residues 64-80 are well-resolved but do not have clear secondary structures. These residues are resolved likely due to their extensive interactions to the intracellular surface of the MBOAT fold (Figure S7A-E). We asked whether the interactions between the N-terminus and the MBOAT fold core may affect the enzymatic activity. Deletion of residues 2-64 (ΔN65) slightly reduced V_max_ (563.9 ± 32.5 nmol/mg/min) but has almost no Δ effect on K_M_ (13.9 ± 2.6 µM, Figure 2J). In contrast, deletion of the entire N-terminus to the first residue of TM1 (residues 2-84, ΔN84) abolishes the enzymatic activity (Figure 2J). The N-Δ terminus is not required for dimer formation because Δ (Figure S7F). To narrow down the region of functional significance, we made three additional N75 and N80, and measured their enzymatic activities. All Δ three have significantly lower enzymatic activity than the wild type, and it appears that larger the deletion, lower the enzymatic activity (Figures 2J and S7G, Table S2). These results are consistent with the recognized role of the N-terminus, however, further structural and functional studies are required to determine the precise impact of the N-terminus on structure and function of hDGAT1.

## The reaction chamber and oleoyl CoA binding site

The MBOAT fold in hDGAT1 carves out a large hollow chamber in the hydrophobic core of the membrane (Figure 3A-D). The almost universally conserved histidine 415 in the MBOAT family of enzymies are found inside of the reaction chamber and on TM7 (Figures 2F and S8). TM2-9 segregates into three groups that form three sidewalls of the chamber: TM2, 3 and 4 pack into a bundle that forms the first sidewall; TM5 and 6 are both very long with almost 40 amino acids each, and the two helices coil into a unit that tilts roughly 56 degree to the membrane norm to form the second sidewall; TM7, 8 and 9 form a panel and the third sidewall (Figure 3A-D). The cytosolic ends of TM7 and 8 is ∼19 Å apart, creating a side entrance to the reaction chamber (Figures 3B and 3E). IL1 and IL2 are located at roughly the cytosolic surface of the membrane and form the floor of the chamber. IL1 (residues 222 to 261) is composed of a helix flanked by two long strands, while IL2 (residues 352 to 396) has a long amphipathic helix (AH, residues 380 to 394) preceded by a short helix and a loop.

**Figure 3.**
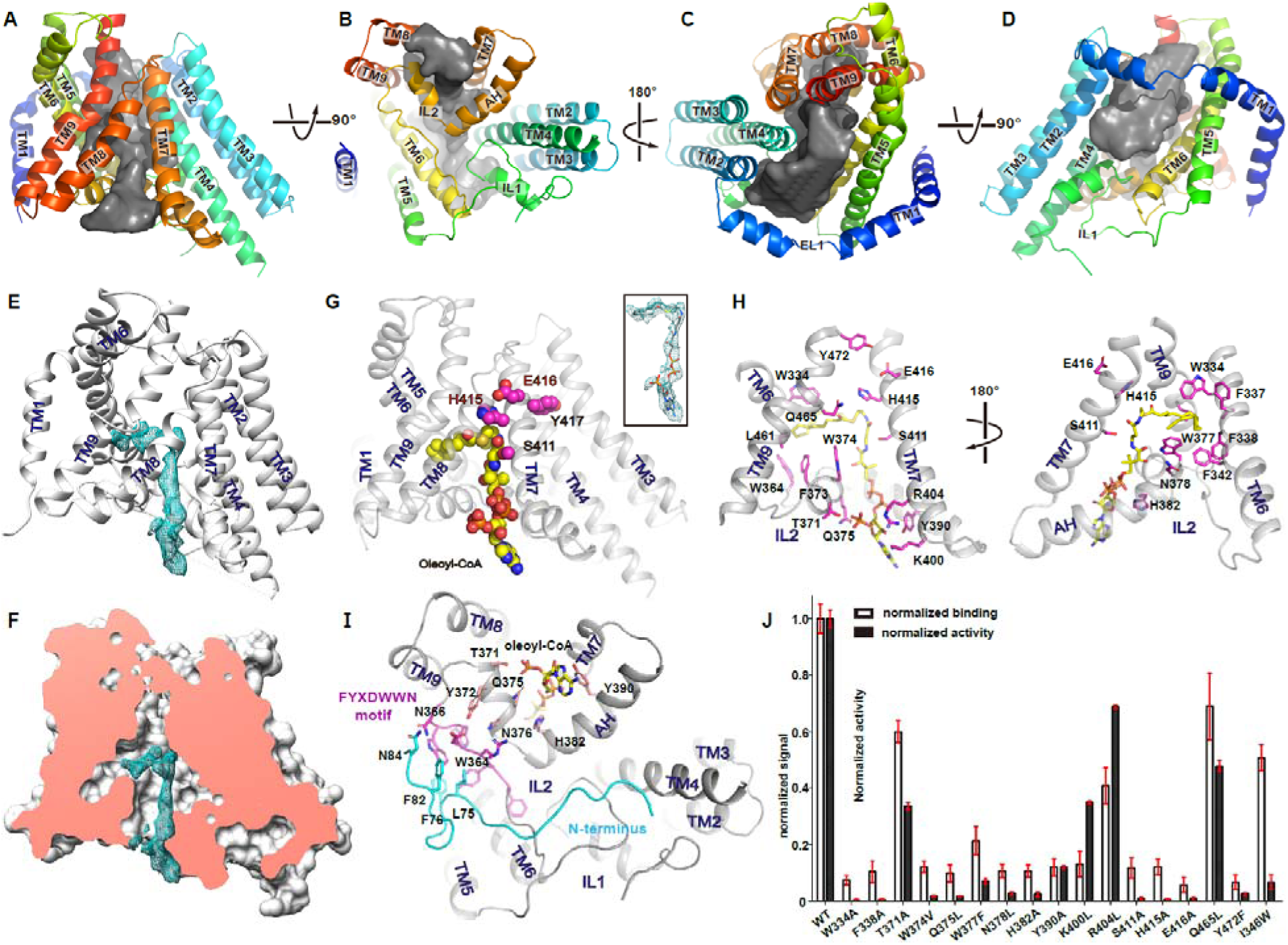
The reaction chamber and oleoyl-CoA binding. **A-D.** The reaction chamber (grey surface) is shown in four orientations with the helices shown as cartoon. **E-F.** Density map of oleoyl-CoA contoured at 7 σ in one hDGAT1 protomer shown as cartoon (**E**) or surface (**F**). In (**G**), an oleoyl-CoA is modeled into the density and shown as spheres with carbon atoms colored in yellow. The conserved SXXHEY motif was labeled as magenta spheres. **H.** Residues that interact with oleoyl-CoA are shown in sticks with carbon atoms colored magenta. **I.** Interaction between the FYXDWWN motif (magenta) with the N-terminus of the neighboring protomer (cyan). **J.** Normalized enzymatic activity and oleoyl-CoA binding of hDGAT1 wild type and mutants. (100 μM of oleoyl-CoA and 200 μM of 1,2-dioleoyl-sn-glycerol). Each bar represents the average of three repeats. Error bars are s.e.m.

The structure of hDGAT1 was solved in the presence of 1mM oleoyl-CoA. A large non-protein density is found at the cytosolic side of the reaction chamber close to IL2 and it extends deep into the reaction chamber (Figures 3E-3F). An oleoyl-CoA can be modeled into this density, with the adenosine 3’,5’-diphosphate at the cytosolic entrance, the 4-phosphate panthothenic acid, β-mercapto-ethylamine extend progressively into the reaction chamber, and the acyl chain residing in a hydrophobic pocket inside of the reaction chamber (Figure 3G). Tyr390, and Lys400 line the entrance of the acyl-CoA binding site, Gln375, Trp377, Asn378, His382 and Ser411 line the tunnel leading to the active site, and Trp334, Phe337, Phe338, Phe342, Trp364, Phe373, Trp374, and Trp377 line the hydrophobic pocket for the acyl chain (Figures 3H and S8A-S8G). The activated thioesther is located to the vicinity of His415, poised for an attack from the activated hydroxyl of DAG. The position of the thioester is stabilizd by interaction between the carbonyl oxygen of the fatty acid and the side chain of Gln465 on TM9 (Figures 3H and S8G). A conserved Pro466 creates a kink on TM9 that brings Gln465 closer to the acyl-CoA (Figure 3H).

IL2 has a crucial role in acyl-CoA binding. Its V-shaped helix-turn-helix motif forms the entrance for the acyl-CoA, and a number of residues on the two helices make direct contact to the acyl-CoA (Figure 3I). The loop preceding the helices contains the FYXDWWN motif, which was identified in both DGAT1 and ACAT as important for enzymatic activites (16, 40, 41) and is highly conserved (Figure S6). Trp364, the first trptophan in the motif, forms part of the hydrophobic pocket for the acyl chain, and although the rest of the motif does not have direct contact with the acyl-CoA, the FYXDWWN motif packs tightly against the helix-turn-helix motif (Figure 3I), and thus mutations in the former could affect the enzymatic activity.

Interestingly, the FYXDWWN motif also makes extensive contact with the N-terminus from the neighboring protomer (Figure 3I), and perturbations to these interactions caused by N-terminal deletions could affect the enzymatic activity although the N-terminus does not make direct contact with the bound oleoyl-CoA.

To assess the functional impact of residues in the active site and ones that line the acyl-CoA binding site, we mutated these residues, one at a time, and measured their enzymatic activity. His415Ala abolishes the enzymatic activity, consistent with its role in catalysis. Point mutations to residues that line the entrance of the acyl-CoA binding site reduces the enzymetic activity by 30 to 70%, while mutations to the rest of the binding pocket, Trp377, Asn378, His382, Ser411, have a larger impact with a loss of more than 80% activity (Figure 3J). Since His415 and Ser411 are part of the highly conserved SxxxHEY motif that was shown to be crucial for enzymatic activity in a related MBOAT enzyme, ACAT (16), we mutated the glutamate and found that Glu416Leu abolishes the enzymatic activities (Figures 3B, 3C and 3E). Although results form these initial mutational studies are largely confirmatory, the structure provides a framework for further studies that will lead to more precise understanding of substrate recognition and the mechanism of catalysis.

## Gateway for DAG and TG

The reaction chamber has a very large opening to the hydrophobic core of the membrane, and the opening is framed by TM4 on one side and TM6 on the other side, and by part of the IL1 (residues 234-245) on the cytosolic side (Figures 3B-D). Residues line the two sides of the entrance are mostly hydrophobic, Val192, Leu196, Met199, and ILE203 on TM4, and Phe337, Leu341, Leu346, and Val349 on TM6 (Figure 4A). A tubular density is found near the entrance and extends into the reaction chamber (Figures 4B and 4C). Although an acyl chain could be modeled into the density, we cannot identify the ligand. We speculate that this large opening would allow entrance of DAG to the reaction chamber from either leaflet of the lipid bilayer, and exit of the product TG (Figure 5). Consistent with this hypothesis, mutating Leu346 to a bulkier side chain Trp produces an enzyme that has no enzymatic activity but retains binding to acetyl CoA (Figure S8I, Table S2).

**Figure 4.**
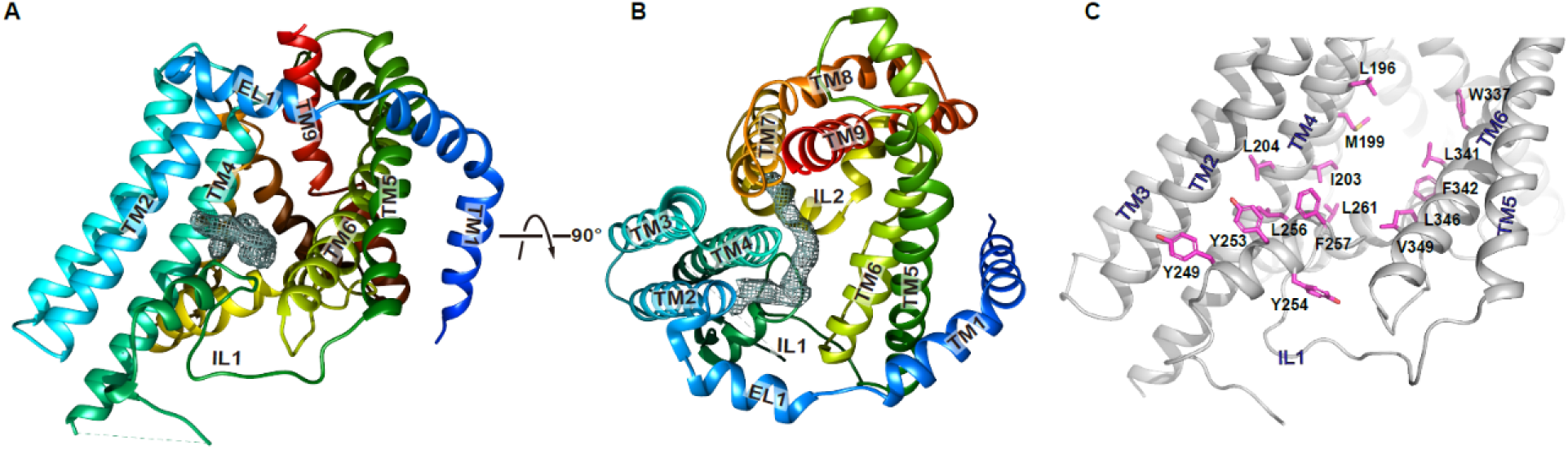
Proposed gateway for DAG entry. **A-B.** An un-modeled tubular density extending from the opening between TM4 and TM5 into the reaction chamber is viewed in two orientations. **C**. Residues that line the TM4-5 opening are labeled as magenta sticks.

**Figure 5.**
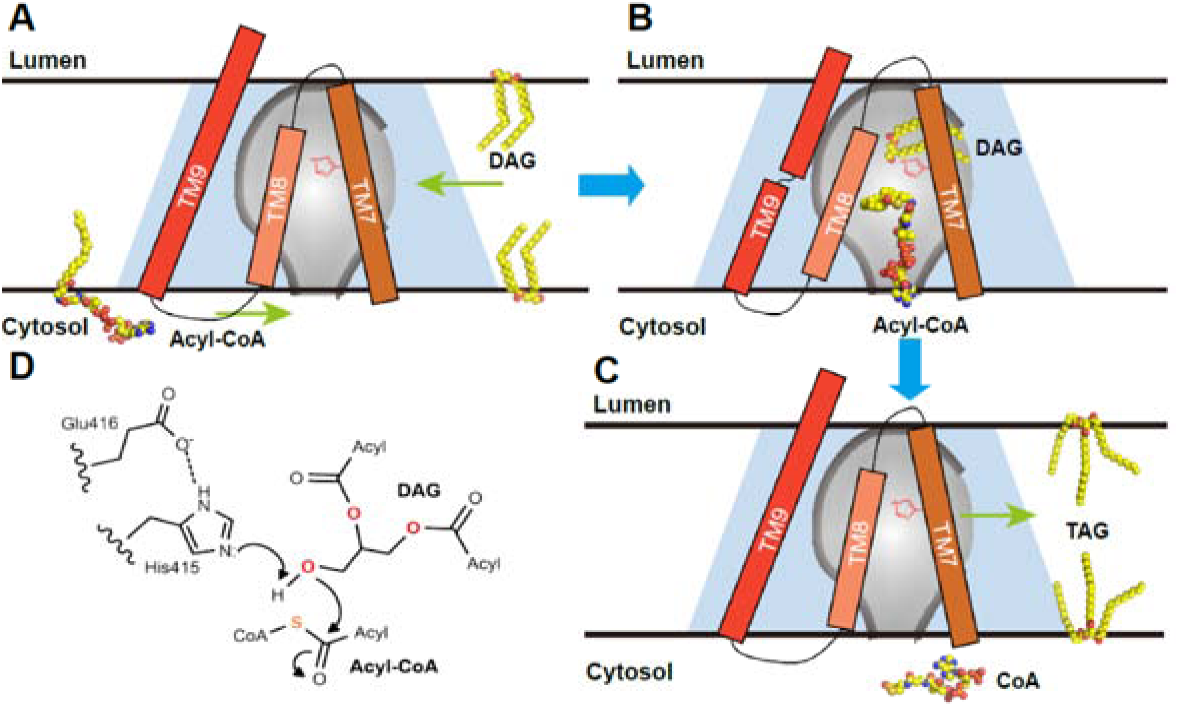
Proposed catalytic mechanism of hDGAT1. **A.** and **B.** A hDGAT1 monomer is shown as a trapezoid in light blue and the reaction chamber in the shape of an inverted flask colored in grey. TM7-9, acyl-CoA and DAG are shown schematically. The catalytic His415 is marked in red on TM7. The CoA moiety of an acyl-CoA binds to hDGAT1 at the cytosolic entrance of the tunnel and the rest of the acyl-CoA slides into the reaction chamber through a slit between TM7 and TM8. The glycerol backbone of a DAG enters the chamber from a side entrance and becomes almost horizontal with the two acyl-chains partially hosted in the hydrophobic core of the membrane. **C.** After the reaction, CoASH exits the chamber through the tunnel and the product TG could diffuse to either leaflet of the membrane. **D.** Proposed catalytic mechanism. Both the 3-hydroxyl of DAG and the thioester of acyl-CoA are positioned near the catalytic H415. E416 helps H415 activate the 3-hydroxyl on DAG for a nucleophilic attack on the thioester of an Acyl-CoA.

## Discussion

hDGAT1 structure defines a conserved MBOAT structural fold, which forms a large chamber in the hydrophobic core of the membrane so that the acyl transfer reaction is isolated inside of the chamber. The structure shows that an acyl CoA is recognized by residues at the cytosolic side of the reaction chamber and a hydrophobic pocket inside of the chamber. Since the hydrophobic acyl chain of an acyl CoA likely is buried in the membrane, we speculate that a slit between TM7 and 8 allows entry of the acyl chain into the chamber (Figure 5A). TM8 is less well resolved than the neighboring helices (Figure S4), suggesting that it is more mobile in the membrane and thus could move to accommodate entry of the acyl chain. DAG likely enters the reaction chamber through the large opening between TM4 and TM6 (Figure 5A). The glycerol backbone of a DAG can reach the catalytic center His415 by interacting with hydrophilic residues in the vicinity of His415 (Figure 5B), while the two hydrophobic aliphatic acyl chains of DAG could remain partially accommodated by the hydrophobic core of the membrane. We speculate that the conserved His415 facilitates the acyl transfer reaction by activating the free hydroxyl on DAG, and the presence of Glu416 could enhance the activation. The activated hydroxyl oxygen then attacks the thioester on the fatty acyl-CoA to form a new ester bond (Figure 5C). The product, TG, could retraces the entrance pathway of DAG back into the membrane while CoA dissipates into the cytosol (Figure 5D).

We noticed that the bacterial DltB protein also has a core of 8 helices that fold into a similar architecture (Figure S9D-F), although it has a total of 11 transmembrane helices (25). In DltB, the intracellular loops are placed more towards the center of the membrane perhaps to accommodate the two substrates coming from either side of the membrane. As a result, the MBOAT fold in DltB does not carve out a reaction chamber in the membrane. The large opening between TM4 and TM6 is covered by two extra helices that are not part of the MBOAT fold (Figure S9A-F). It is interesting to note that the binding site for acyl-CoA in hDGAT mirrors the substrate binding site in DltB. DltC, with a covalently linked D-alanyl Ppant group, is a substrate to DltB and equivalent to a fatty acyl-CoA to hDGAT1. In the DltB-DltC complex, DltC is positioned at the equivalent location of acyl-CoA to the MBOAT fold. Overall, DltB is shaped like an hourglass that allows the two substrates to approach the reaction center from either sides of the membrane, and the transfer of an acyl group across the membrane (Figure S9G-J). These observations highlight the versatility of the MBOAT fold that can be adapted for different functions.

## Acknowledgments

This work was supported by grants from NIH (DK122784 and HL086392 to MZ), Cancer Prevention and Research Institute of Texas (R1223 to MZ), the Robert Welch Foundation (Q1279 to BVVP), Ara Parseghian Medical Research Foundation (to N.Y. and Y.H.), and the New Jersey Council for Cancer Research (to H.Q.). N.Y. is supported by the Shirley M. Tilghman endowed professorship from Princeton University. We thank Paul Shao for technical support during EM image acquisition. We acknowledge the use of Princeton’s Imaging and Analysis Center, which is partially supported by the Princeton Center for Complex Materials, and the National Science Foundation (NSF)-MRSEC program (DMR-1420541).

## Author Contributions

M.Z., L.W., Y.N. and Z.R. conceived the project. L.W., Y.N., H.Q., Z.R., Y.H., H.Z. conducted experiments. L.W., Y.N., H.Q., Z.R., Y.H., N.Y., and M.Z. analyzed data. L.H. and B.V.V.P. advised on model building and refinement. L.W., Z.R. and M.Z. wrote the initial draft and all authors participated in revising the manuscript.

## Methods

### Cloning, expression, and purification of human DGAT1

Human DGAT1 gene (accession number NP_036211) was codon-optimized and cloned into a pFastBac dual vector(42) for production of baculovirus by the Bac-to-Bac method (Invitrogen). High Five Cells (Thermofisher) at a density of ∼3×10^6^ cells/ml were infected with baculovirus and grown at 27 °C for 48–56 h before harvesting. Cell membranes were prepared following a – previous protocol (42) and frozen in liquid nitrogen.

Purified membranes were thawed and homogenized in 20 mM HEPES, pH 7.5, 150 mM NaCl and 2mM β-mercaptoethanol, and then solubilized with 1% (w/v) Lauryl Maltose Neopentyl β Glycol (LMNG, Anatrace) at 4 °C for 2 h. After centrifugation (55,000g, 45min, 4 °C), hDGAT1 was purified from the supernatant using a cobalt-based affinity resin (Talon, Clontech) and the His-tag was cleaved by TEV protease. Oleoyl-CoA (20 μM) was added to reduce aggregation, and hDGAT1 was then concentrated to 5 mg/ml (Amicon 100 kDa cutoff, Millipore) and loaded onto a size-exclusion column (SRT-3C SEC-300, Sepax Technologies, Inc.) equilibrated with 20 mM HEPES, pH7.5, 150 mM NaCl, 0.01% glyco-diosgenin (GDN, Anatrace) for cryo-EM grid preparation. For enzymatic assays, GDN was replaced with 1 mM (w/v) n-dodecyl--D-βmaltoside (DDM, Anatrace).

hDGAT1 mutants were generated using the QuikChange method and the entire cDNA was sequenced to verify the mutation. Mutants were expressed and purified following the same protocol as wild type.

### Cryo-EM sample preparation and data collection

The cryo grids were prepared using Thermo Fisher Vitrobot Mark IV. The Quantifoil R1.2/1.3 Cu grids were glow-discharged with air for 40 sec at medium level in a Plasma Cleaner (Harrick Plasma, PDC-32G-2). Purified hDGAT1 was mixed with 1 mM of oleoyl-CoA and concentrated to approximately 20 mg/ml. Aliquots of 3.5 µl purified hDGAT1 were applied to glow-discharged grids. After being blotted with filter paper (Ted Pella, Inc.) for 3.5 s, the grids were plunged into liquid ethane cooled with liquid nitrogen. A total of 2706 micrograph stacks were collected with SerialEM(43) on a Titan Krios at 300 kV equipped with a K2 Summit direct electron detector (Gatan), a Quantum energy filter (Gatan) and a Cs corrector (Thermo Fisher), at a nominal magnification of 105,000 × and defocus values from -2.0 µm to -1.2 µm. Each stack was exposed in the super-resolution mode for 5.6 s with an exposing time of 0.175 s per frame, resulting in 32 frames per stack. The total dose rate was about 50 e^-^/Å^2^ for each stack. The stacks were motion corrected with MotionCor2 (44) and binned 2 fold, resulting in a pixel size of 1.114 Å/pixel. In the meantime, dose weighting was performed (45). The defocus values were estimated with Gctf (46).

### Cryo-EM data processing

A total of 2,749,110 particles were automatically picked with RELION 2.1 (47–49). After 2D classification, a total of 1,000,063 particles were selected and subject to a guided multi-reference classification procedure. The references, one good and three bad, were generated with limited particles in advance (Figure S3). Particles selected from multi-references 3D classification were subjected to a global angular search 3D classification with one class and 40 iterations. The outputs of the 31th-40th iterations were subjected to local angular search 3D classification with four classes separately. Particles from the good classes of the local angular search 3D classification were combined, yielding a total of 408945 particles. After handedness correction and C2 symmetry application, 3D auto-refinement with an adapted mask yielded a reconstruction with an overall resolution of 3.1 Å. Further 3D classification yielded a class of 275,945 particles and after 3D auto-refinement, yielded a map of 3.1 Å with improved density of TM2, TM3, TM8 and lipids.

All 2D classification, 3D classification, and 3D auto-refinement were performed with RELION 3.0 and Cryosparc (50). Resolutions were estimated with the gold-standard Fourier shell correlation 0.143 criterion (51) with high-resolution noise substitution (52).

### Model building and refinement

For *de novo* model building of hDGAT1, a ploy-Alanine model was first built into the 3.1Å density map manually in COOT (53). Structure refinements were carried out by PHENIX in real space with secondary structure and geometry restraints (54). The EMRinger Score was calculated as described (55).

### DGAT1 Activity assay

hDGAT1 activity was monitored using a fluorescence-based coupled-enzyme assay (56) in a quartz cuvette at 37°C. The cuvette was read in a FluoroMax-4 spectrofluorometer (HORIBA) with 340 nm excitation and 465 nm emission at 15 s internals. All assays were done in a buffer with 20 mM HEPES, pH 7.5, 150 mM NaCl, 2 mM β-mercaptoethanol, 0.5 mM DDM, 1% TritonX-100. Final concentrations of NAD^+^, thiamine pyrophosphate and α-ketoglutarate were 0.25 mM, 0.2 mM and 2 mM respectively. The α-ketoglutarate dehydrogenase (αKDH) was prepared from beef heart using a published protocol (57). Sufficient amount of αKDH was used to ensure that the hDGAT1 reaction is the rate limiting step. The hDGAT1 concentration in the assay was 40 nM. The oleoyl-CoA concentration was 2.5-100 µM in assays for K_M_ and V_max_ determination, and 100 µM in all other tests. The concentrations of 1,2-dioleoyl-sn-glycerol and other monoacylglycerols were 200 µM in the selectivity assay. The initial rates in various DG concentrations were not well fit with the traditional Michaelis Menten equation, but could be fit with an allosteric sigmoidal equation: Y=Vmax*X^h^/(Km+X^h^), in which X is DAG concentrations, and h is the Hill coefficient.

### Scintillation proximity assay

Binding of Oleoyl-CoA to hDGAT1 was estimated using a Scintillation Proximity Assay (SPA). Purified hDGAT1 (with his tag) was absorbed onto Copper HIS-Tag PVT beads (Perkin Elmer, RPNQ0095) and incubated with [^3^H]-Acetyl-CoA (ARC, ART0213A) for 30 min at ∼22°C in the binding buffer (20 mM Hepes pH 7.5, 150 mM NaCl, and 0.02% GDN). Each 100 µL reaction mixture contains 600 ng hDGAT1, 0.25 µM [^3^H]-Acetyl-CoA and 2.5 mg/ml Copper HIS-Tag PVT beads. Background binding was estimated in the presence of both 800 mM imidazole or in the absence of protein. For dose response curves, 0.005 to 200 µM of cold oleoyl-CoA was used to compete for the binding of 0.25 µM 3H-acetyl-CoA. The binding assay was performed in a 96-well plate (Perkin Elmer) and the scintillation read out by a MicroBeta 2450 Microplate Counter (Perkin Elmer). Data were plotted in Graphpad 8.0 software and fit with the following equation to obtain IC_50_: Y=Bottom+(Top-Bottom)/(1+10^(X-LogIC_50_))

## Supplemental Information

**Figure S1.**
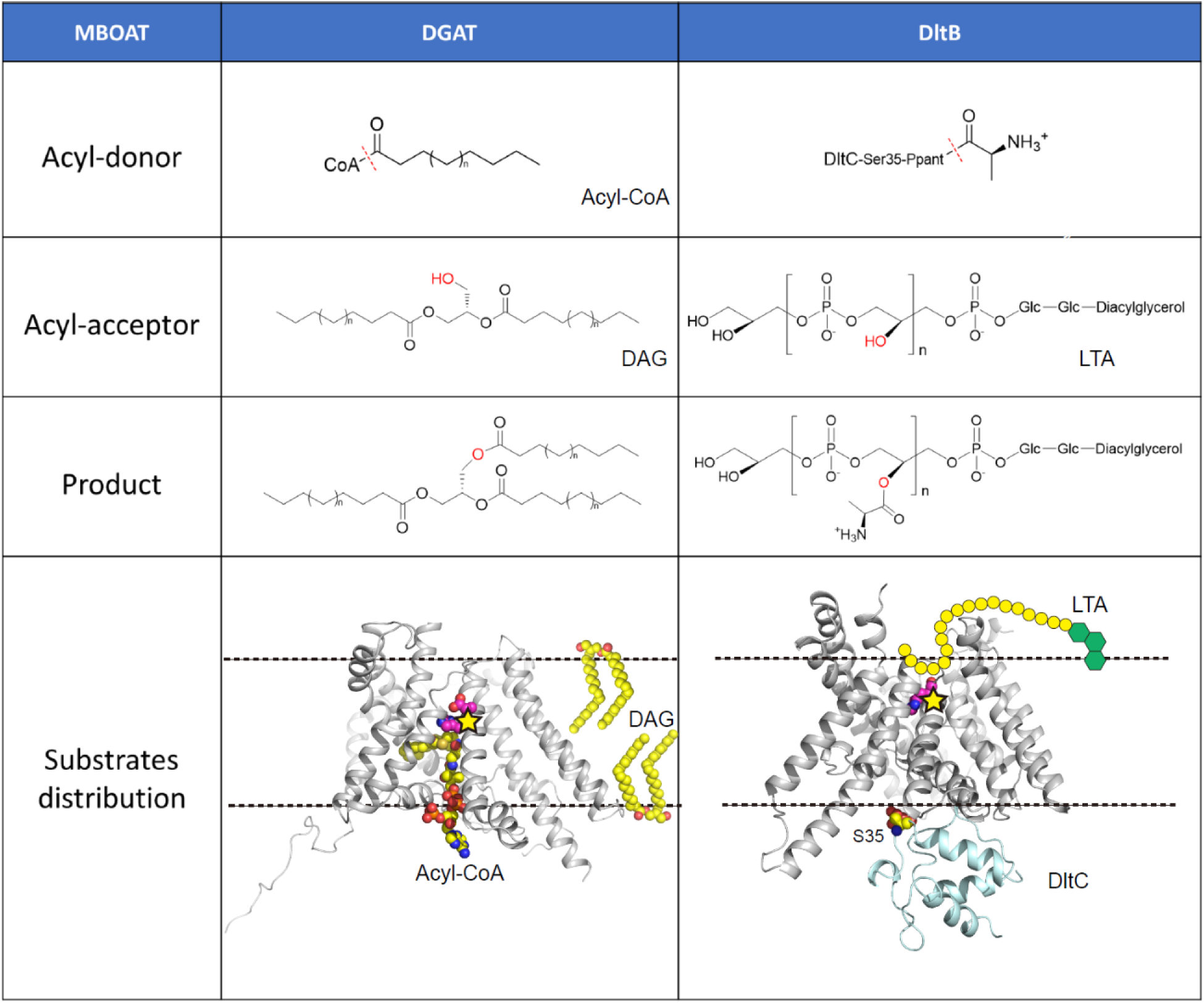
Comparison DltB and DGAT1. Both hDGAT1 and DltB have an acyl-group donor and an acceptor. In the acyl-group donor row, the red dashed lines indicate the bonds that are broken during acyl-transfer reactions. In the acyl-group acceptor row, the hydroxyl groups are highlighted in red. In hDGAT1, acyl-CoA comes from the intracellular side while DAG comes from inside of the membrane. In DltB, the Ppant-DltC is intracellular while the LTA is extracellular.

**Figure S2.**
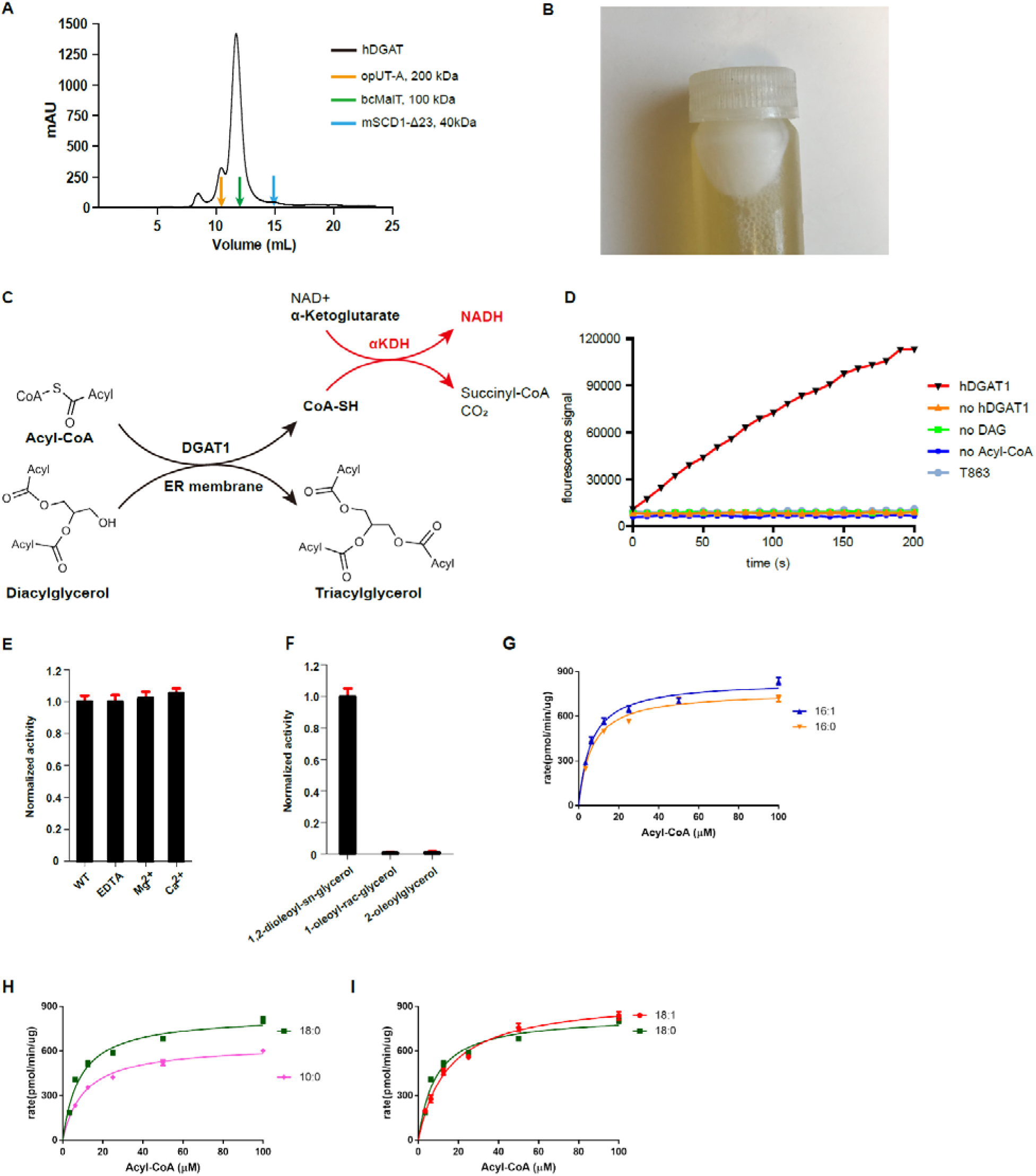
Biochemical and functional characterization of hDGAT1. **A.** size exclusion profile of hDGAT1. Elution volume of membrane proteins of known molecular weight, bcMalT (100 kDa, green) (58), mouse SCD1 (41 kDa, blue) (42) and opossum UT-A (200 kDa) (59) are marked by arrows. **B.** A white layer of fat appeared after membrane solubilization and centrifugation. **C.** hDGAT1 reaction is coupled to that of α-ketoglutarate dehydrogenase (αKDH) to monitor production of Coenzyme A in real time. **D.** Fluorescence of NADH plotted versus M oleoyl-CoA in the presence of αKDH, NAD^+^ (0.25 mM), α-ketoglutarate (2 mM), and thiamine pyrophosphate (0.2 mM). **E.** Normalized activity of hDGAT1 in the presence of EDTA, 1mM of Ca^2+^ or Mg^2+^. Each reaction has 100 μM of oleoylCoA and 200 M of 1,2-dioleoyl-sn-glycerol. **F**. Normalized activity of hDGAT1 in the presence of 200 μM of 1,2-dioleoyl-sn-glycerol, 1-oleoyl-sn-glycerol, or 2-oleoyl-sn-glycerol. Each reaction has 100 μM of oleoyl-CoA. **G-I**. Enzymatic activity of hDGAT1 with different acyl-CoAs. Solid line represents fit of the data points with a Michaelis-Menten equation. In **E**-**H**, each symbol represents the average of three repeats. Error bars are s.e.m.

**Figure S3.**
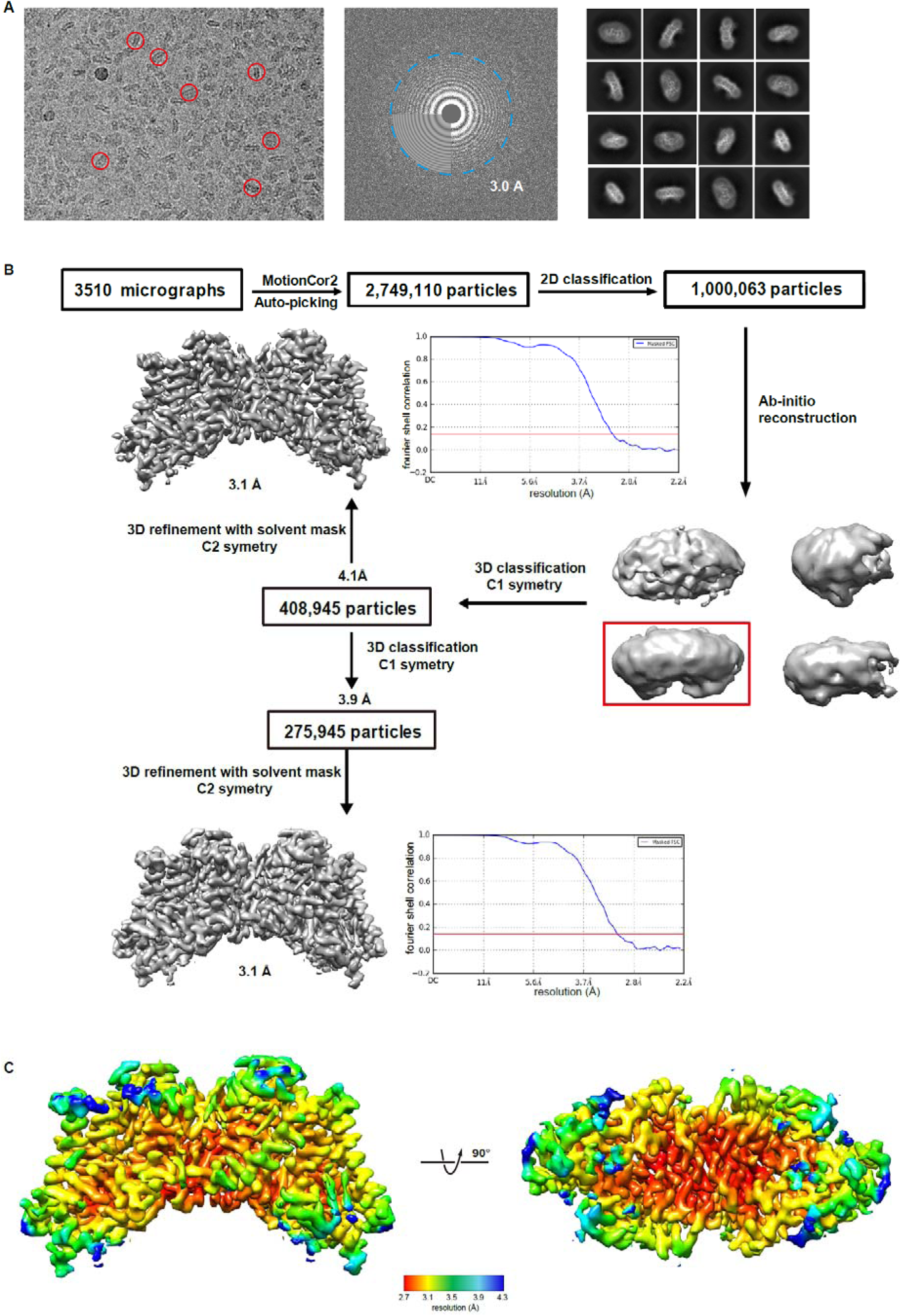
Cryo-EM Analysis of the hDGA1. **A.** Representative micrograph of hDGAT1 (left), its Fourier transform (middle) and representative 2d class averages (right). Representative particles are labeled in red circles. **B.** A flowchart for the cryo-EM data processing and structure determination of the hDGAT1. Final maps of hDGAT1 and the gold-standard Fourier shell correlation curves for the overall maps are shown. **C**. Local resolution maps calculated using RELION 2.0.

**Figure S4.**
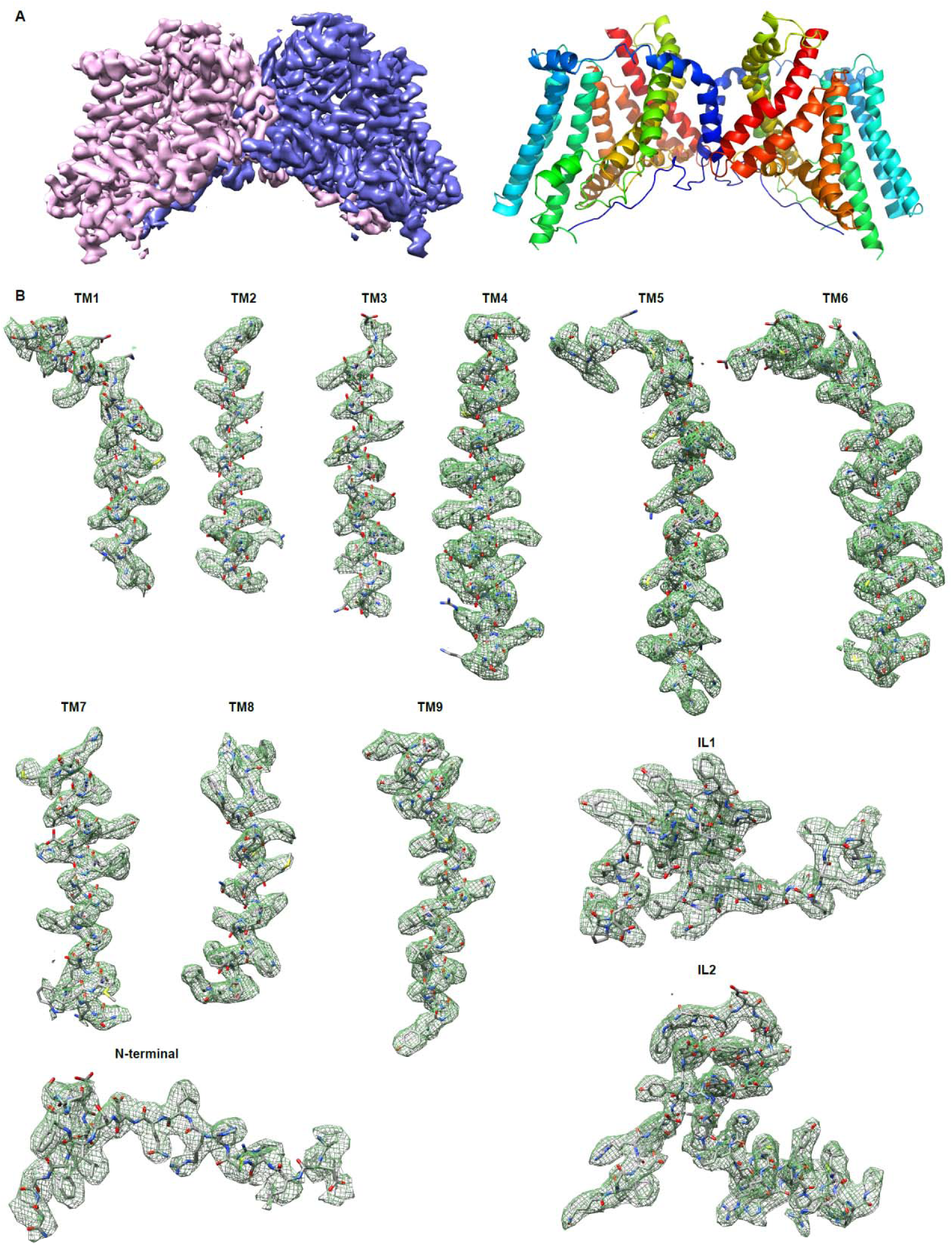
EM Maps for hDGAT1. **A.** The overall map of hDGAT1 and its atomic model. **B.** EM density for each TM helix, each intracellular loop and the N-terminus.

**Figure S5.**
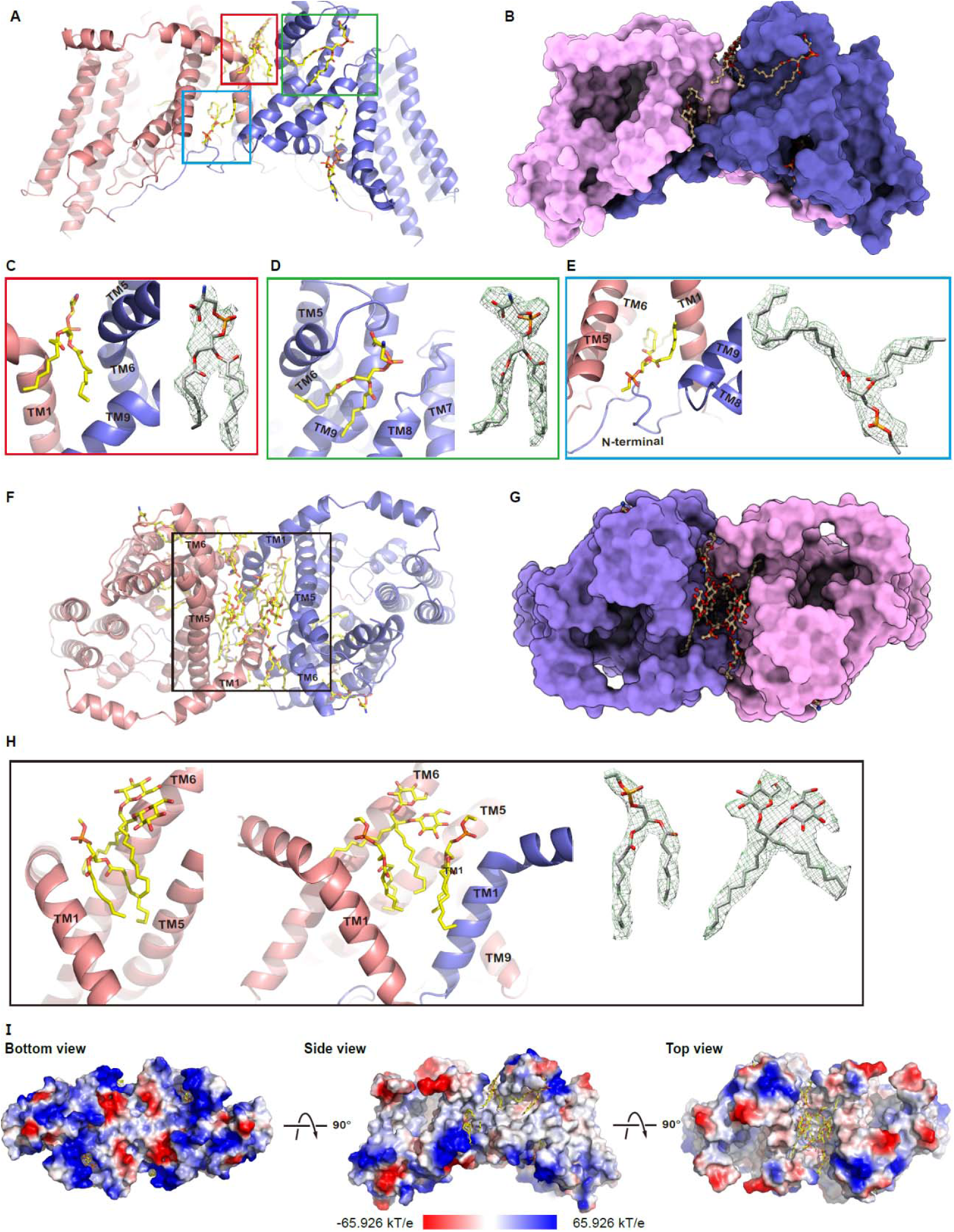
Dimerization interface and lipids. **A-H.** hDGAT1 dimer is shown in two orientations as cartoon (**A** and **F**) and surface (**B** and **G**) representation. Lipids and detergent molecules buried in the dimer interface or attached to the surface of hDGAT1 are shown as sticks. **C**-**E and H**. Detailed view of each detergent/lipid molecule and its corresponding density. **I**. hDGAT1 dimer is shown in three orientations as electrostatic surface representation. The electrostatic potential is calculated using the APBS plugin (60).

**Figure S6.**
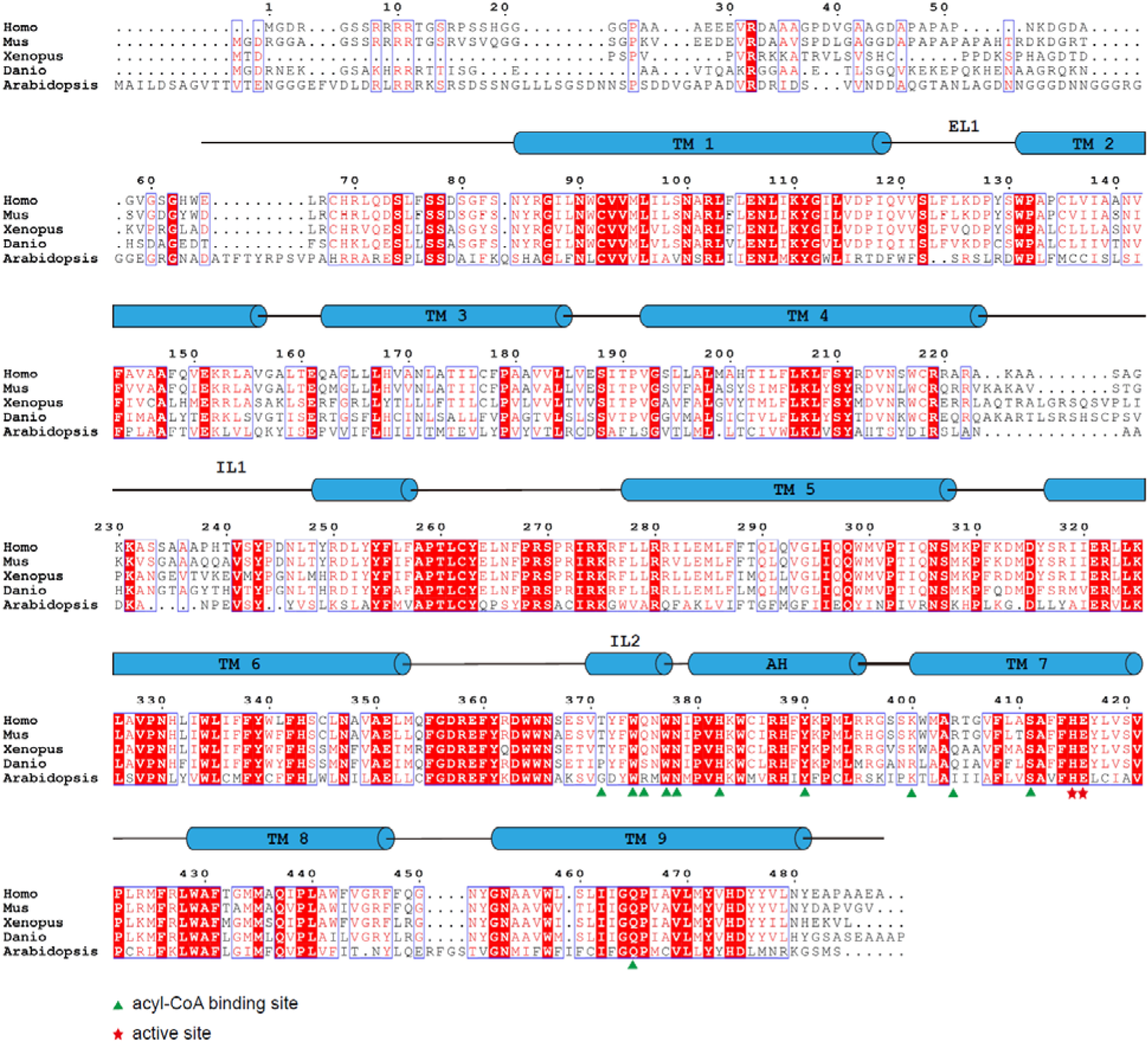
DGAT1 sequence alignment. DGAT1 sequences of human (Uniprot accession number O75907), mouse (Q9Z2A7), frog (XENLA, A0A1L8G0L4), fish (DANIO, Q6P3J0), thale cress (ARABID, Q9SLD2) and human ACAT (P35610) are aligned using the Clustal Omega server (61). Secondary structural elements of hDGAT1 are labeled above the alignment. Residues are colored based on their conservation using the ESPript server (62). Residues at the acyl-CoA binding site are labeled with green triangles and those at the active site with red stars.

**Figure S7.**
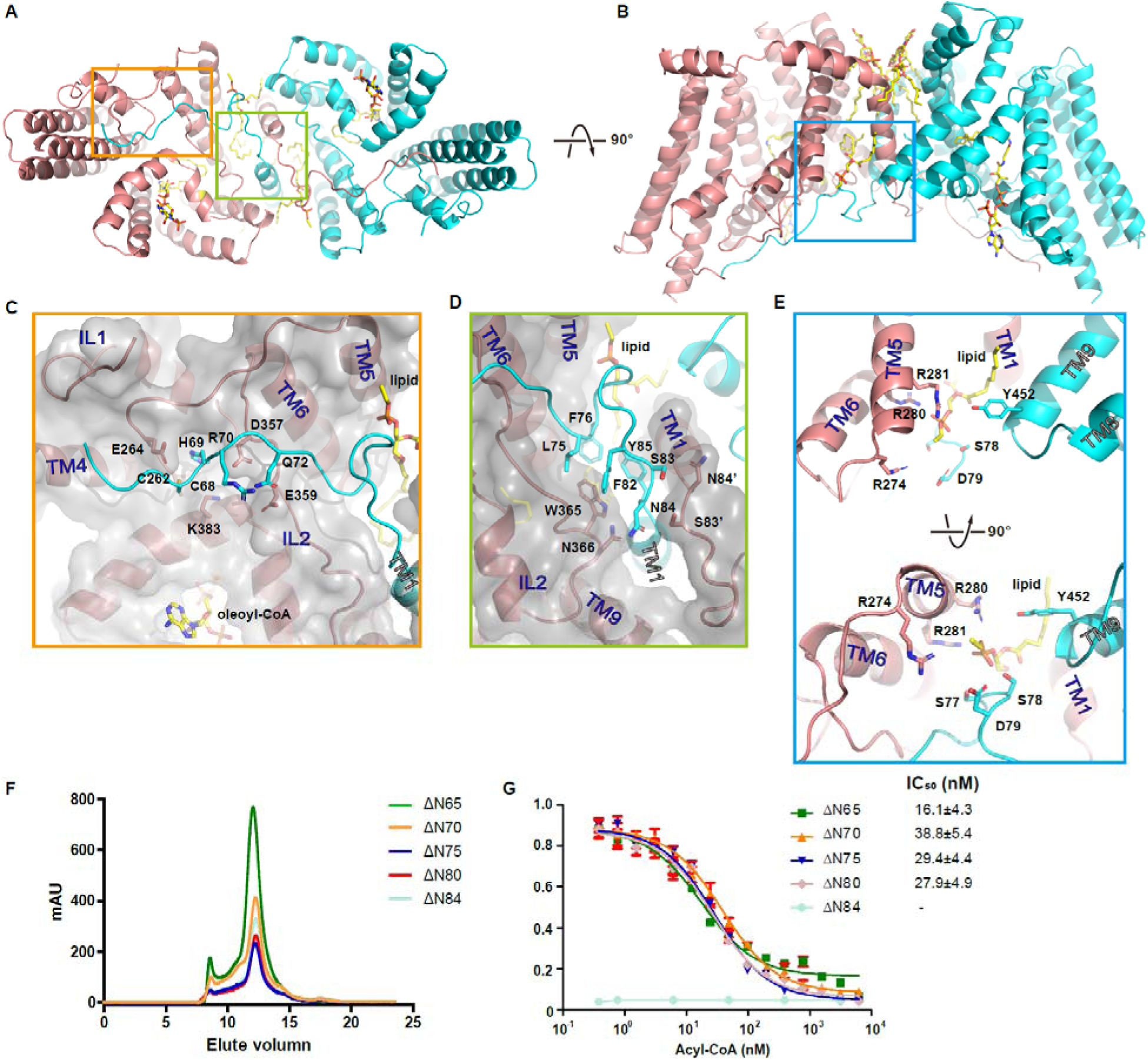
Interaction of the N-terminus with the neighboring protomer. **A.** hDGAT1 dimer (cartoon) is viewed in two orientations. Detailed interactions between a hDGAT1 protomer and the N-terminus from the neighboring protomer are shown in **C-E**. Residues involved in the interactions are shown as sticks. **F**. Size-exclusion chromatography of N-terminal truncations of hDGAT1. **G**. Competitive binding of various concentrations of cold oleoyl-CoA against 0.25 μM of ^3^H-acetyl-CoA on different hDGAT1 N-terminal truncation mutations. Each symbol is the average of three repeats, and error bars are s.e.m..

**Figure S8.**
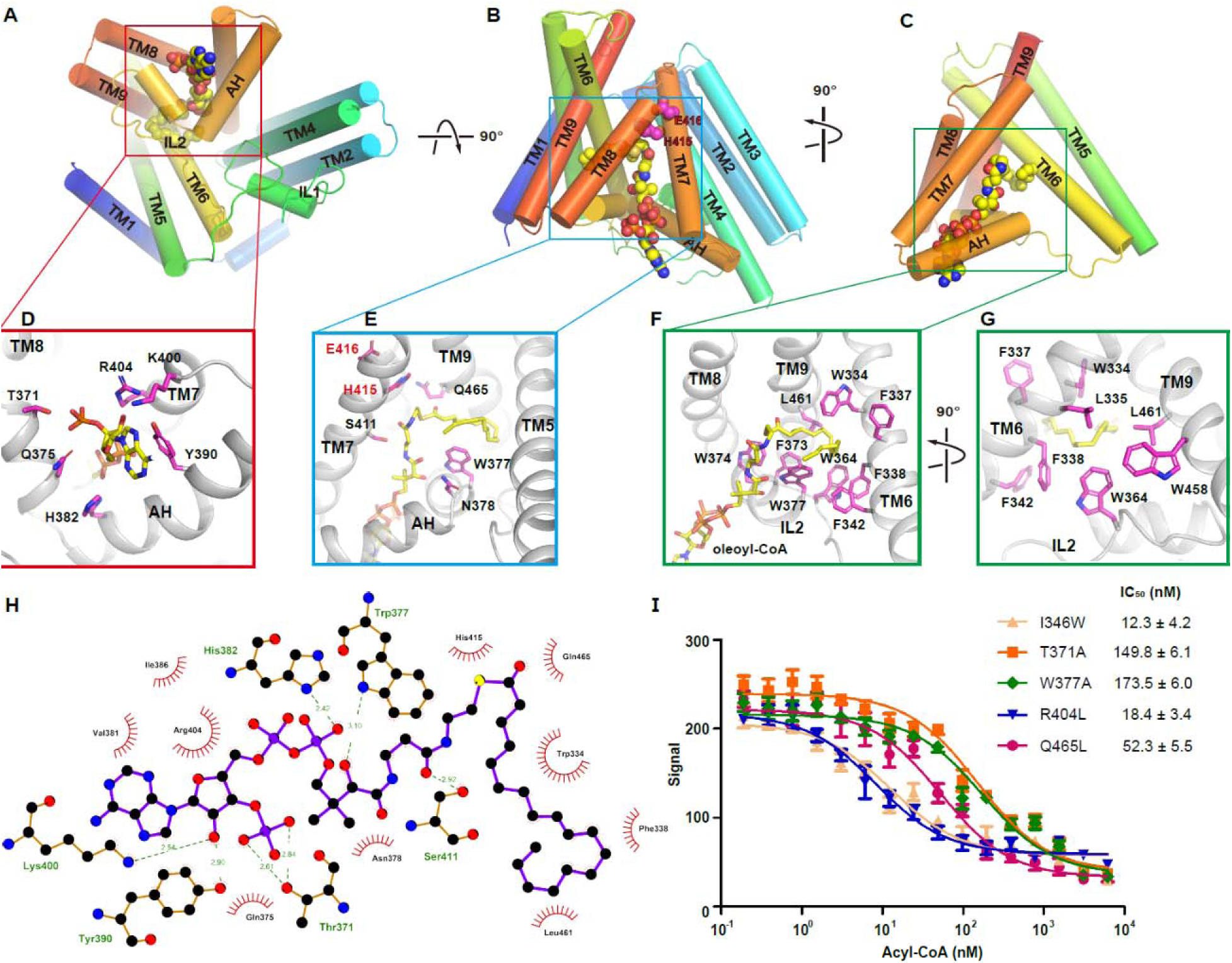
Oleoyl-CoA binding binding site. **A-C.** Oleoyl-CoA (spheres) bound to hDGAT1 protomer (cartoon) viewed in three orientations. Detailed interaction between hDGAT1 and the CoA moiety and the acyl-chain are shown in **D-G**. Residues involved in coordinating oleoyl-CoA are shown as sticks with carbon atoms colored in magenta. **H**. Planar view of the interaction between oleoyl-CoA and hDGAT1 generated by LigPlus (63, 64). **I**. Competitive binding for the residues involved in oleoyl-CoA binding. Each symbol represents the average of three repeats, and error bars are s.e.m.. Data were fit to a single site binding isotherm.

**Figure S9.**
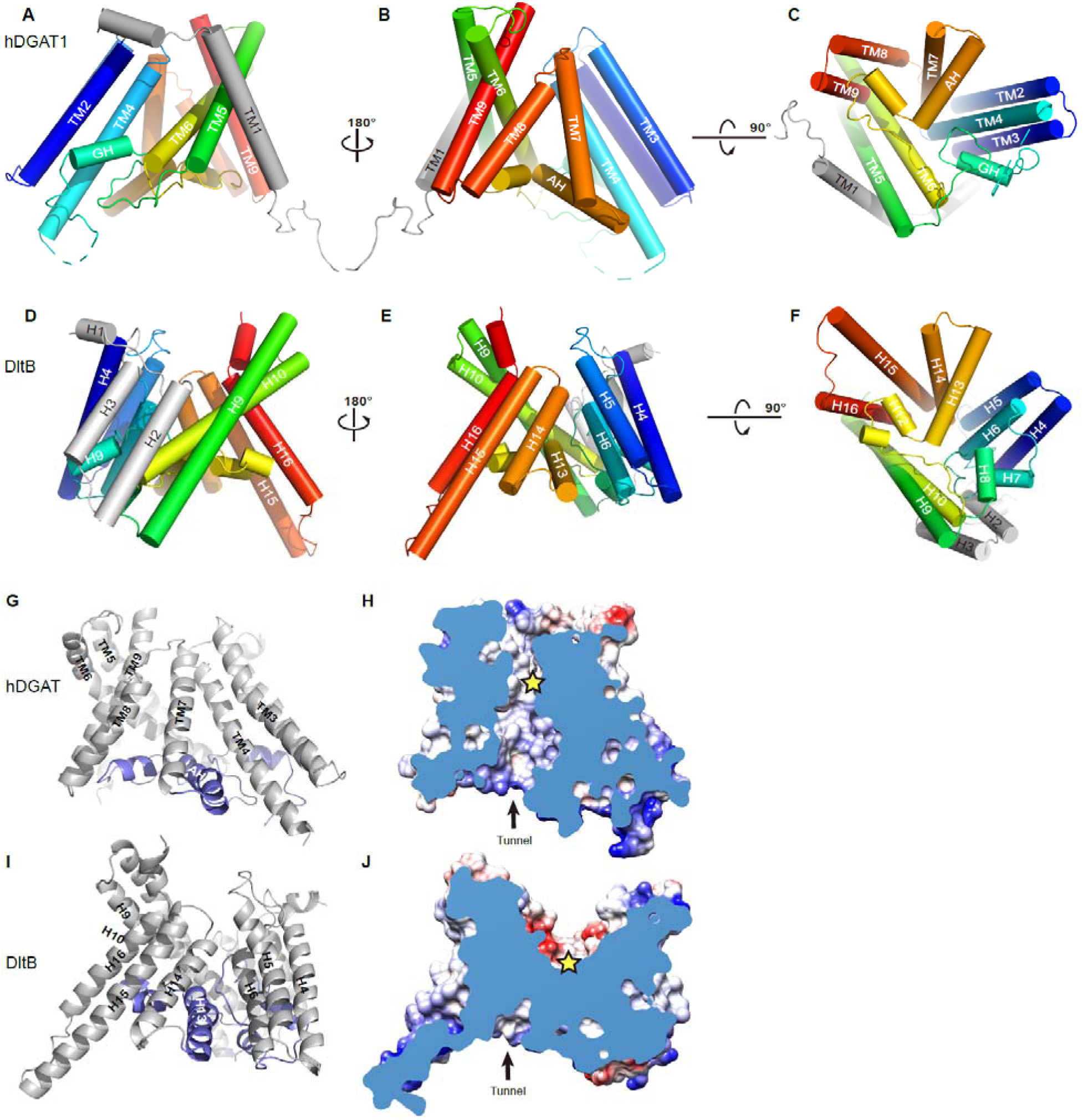
The MBOAT fold in hDGAT1 and DltB. Structures of hDGAT1 (**A-C**) and DltB (**D-F**) are shown as cartoon in three different orientations. The equivalent helices in the MBOAT fold are colored the same. Helices that are distinct in each protein, TM1 in hDGAT1 and H1-H3 in DtlB, are colored in grey. **G** and **I.** Cartoon representations of hDGAT1 and DltB structure. AH2 in hDGAT1 and H13 in DltB adopts very different conformations and are highlighted in blue. **H** and **J.** Cut-away surface illustrations of hDGAT1 and DltB showing their cytosolic tunnels. The position of the conserved histidine residue is marked as a yellow star. In DltB, active site is located to the thin layer separating the intra- and extracellular sides.

**Table S1.** Summary of Cryo-EM data collection, processing and refinement.

| **Protein** | **hDGAT1** |
| --- | --- |
| **Cryo-EM Data Collection** |  |
| Voltage (kV) | 300 |
| Magnification (x) | 105,000 |
| Pixel Size (Å) | 1.114 |
| Electron exposure (e-/Å<sup>2</sup>/frame) | 1.56 |
| Defocus range (µm) | [-2.0, -1.2] |
| Number of image stacks | 3510 |
| Number of frames per stack | 32 |
| **Cryo-EM Data Processing** |  |
| Initial number of particles | 2,749,110 |
| Final number of particles | 275,945 |
| Symmetry imposed | C2 |
| Map sharpening B factor (Å<sup>2</sup>) | -100 |
| Map resolution (Å) | 3.1 |
| Map resolution range (Å) | 3.1-4.3 |
| FSC threshold | 0.143 |
| **Model Refinement** |  |
| Number of amino acids | 808 |
| Total non-hydrogen atoms | 7204 |
| Average B factor (Å<sup>2</sup>) | 85.70 |
| Bond length r.m.s.d. (Å) | 0.015 |
| Bond angle r.m.s.d. (°) | 1.146 |
| Ramachandran Plot |  |
| Favored (%) | 94.76 |
| Allowed (%) | 4.24 |
| Outliers (%) | 0.00 |
| Rotamer outliers (%) | 0.86 |
| EMRinger Score | 3.28 |

**Table S2.** Summary of enzymological parameters of hDGAT1 and mutants.

| substrates/mutants/truncations | $K_m$ (acyl-CoA, $\mu\text{M}$) | $V_{\max}$ (acyl-CoA, nmol/mg/min) | $IC_{50}$ (oleoyl-CoA, nM) |
| --- | --- | --- | --- |
| oleoyl-CoA | $14.6 \pm 1.3$ | $956.6 \pm 36.1$ | $18.2 \pm 5.2$ |
| stearoyl-CoA | $8.6 \pm 1.3$ | $839.4 \pm 49.9$ | - |
| palmitoleoyl-CoA | $6.2 \pm 0.9$ | $838.6 \pm 31.6$ | - |
| palmitoyl-CoA | $6.4 \pm 1.1$ | $767.8 \pm 34.0$ | - |
| octodecyl-CoA | $10.5 \pm 1.4$ | $642.9 \pm 25.0$ | - |
| 1,2-dioleoyl-sn-glycerol | $597.1 \pm 94.5$ | $2335 \pm 279.1$ (h = 1.5) | - |
| I346W | - | - | $12.3 \pm 4.2$ |
| T371A | - | - | $149.8 \pm 6.1$ |
| W377A | - | - | $137.5 \pm 6.0$ |
| R404L | - | - | $18.4 \pm 3.4$ |
| Q465L | - | - | $52.3 \pm 5.5$ |
| $\Delta\text{N}65$ | $13.9 \pm 2.6$ | $563.9 \pm 32.5$ | $16.1 \pm 4.3$ |
| $\Delta\text{N}70$ | $24.4 \pm 1.5$ | $540.7 \pm 25.5$ | $38.8 \pm 5.4$ |
| $\Delta\text{N}75$ | $28.2 \pm 3.1$ | $497.8 \pm 21.9$ | $29.4 \pm 4.4$ |
| $\Delta\text{N}80$ | $28.6 \pm 4.2$ | $276.1 \pm 16.4$ | $27.9 \pm 4.9$ |
| $\Delta\text{N}84$ | - | - | - |

